# Dynamics of inhibitory control during bilingual speech production: An electrophysiological study

**DOI:** 10.1101/596445

**Authors:** Xiaochen Zheng, Ardi Roelofs, Hasan Erkan, Kristin Lemhöfer

**Author notes:** Correspondence to Xiaochen Zheng, Donders Institute for Brain, Cognition and Behaviour, Radboud University, P. O. Box 9104, 6500 HE, Nijmegen, The Netherlands.

## Abstract

Bilingual speakers have to control their languages to avoid interference, which may be achieved by enhancing the target language and/or inhibiting the nontarget language. Previous research has provided evidence that bilinguals may use inhibition (e.g., Jackson, Swainson, Cunnington, & Jackson, 2001), which is reflected in the N2 component of the event-related potential (ERP). In the current study, we investigated the dynamics of inhibitory control by measuring the N2 during language switching and repetition in picture naming. We recorded the EEG of 30 unbalanced Dutch-English bilinguals in a cued language-switching task. Participants had to name pictures in Dutch or English depending on the cue. A run of same-language trials could be short (two or three trials) or long (five or six trials). We assessed whether RTs and N2 changed over the course of same-language runs, and at a switch between languages. Results showed that speakers named pictures more quickly late as compared to early in a run of same-language trials. Moreover, they made a language switch more quickly after a long run than after a short run. In ERPs, we observed a widely distributed switch effect in the N2, which was larger after a short run than after a long run. The N2 was not modulated during a same-language run, challenging Kleinman and Gollan (2018), who maintained that inhibition accumulates over time. Our results suggests that speakers adjust the level of inhibitory control at a language switch, but not when repeatedly naming in the same language.

## Introduction

Bilingual speakers can usually stay in the same language or switch between languages fluently, but this process is not as effortless as it appears to be. In order to properly speak one language and avoid interference from the other, bilinguals need to control their languages in use. This may be achieved by inhibiting the nontarget language and/or enhancing the target language (Allport & Wylie, 1999; Meuter & Allport, 1999). With repeated use of the same language, the target language becomes increasingly activated and speaking becomes easier. Nevertheless, it remains unclear how language control unfolds over time. Is less control required when the target language is fully functioning, due to a high degree of bottom-up activation? Or is (more) top-down control also contributing to the fact that producing the target language gets easier? What are the consequences of having used the same language for a prolonged period of time when speakers have to switch to a different language? In the current study, we investigate the dynamics of language control during language repetition and switching.

Language control is commonly studied using a bilingual picture-naming paradigm, where speakers are asked to name pictures and switch languages according to a given cue (a flag, a color patch, etc). As concerns naming reaction time (RT), bilingual speakers are usually slower when they have to switch to a different language compare to repeatedly naming in the same language, known as *switch cost*. Paradoxically, switching to the dominant first language (L1) is usually more costly than to the weaker second language (L2) (e.g., Gollan, Kleinman, & Wierenga, 2014; Meuter & Allport, 1999). It is hypothesized that during L2 repetition, more top-down control is required to inhibit the dominant L1, or enhancing the weaker L2, compared to L1 repetition. Therefore, it becomes more difficult to overcome the residual inhibition or enhancement when switching back to the L1 than vice versa (Allport & Wylie, 1999; Green, 1998). In a mixed-language context, speaking in the L1 can become more difficult even outside switch trials, and thus become slower compared to the L2 (*reversed dominance effect*; Christoffels, Firk, & Schiller, 2007; Costa & Santesteban, 2004; Verhoef, Roelofs, & Chwilla, 2009; see Declerck & Philipp, 2015, for a more extensive discussion on the relationship between top-down control, asymmetric switch cost, and the reversed dominance effect). Inhibition is considered to be one of the main forces of the language control process (e.g., Jackson, Swainson, Cunnington, & Jackson, 2001; Green, 1998; Verhoef, Roelofs, & Chwilla, 2010). However, evidence seems to diverge on how inhibitory control unfolds over time (Kleinman & Gollan, 2018; Zheng, Roelofs, & Lemhöfer, 2018).

Using the bilingual picture-naming paradigm, Zheng and colleagues (2018) observed that top-down control (i.e., inhibition of the nontarget language and/or enhancement of the target language) decreases with repeated use of the same language. In that study, the number of same-language trials before a switch (i.e., run length) was manipulated. Results showed that bilingual speakers’ responses were slower and less accurate when switching after a short run compared to a long run. This was explained as follows. With repeated use of the same language, the target language becomes more activated and thus less top-down control is needed. As a consequence, it is harder to switch after a short run (when more control is still applied) compared to a long run (when less control is applied), with more residual control to overcome. Interestingly, the *run-length effect* was only present when switching to the L1 rather than to the L2. The difference between languages seems due to the fact that the weak L2 competes less for selection during L1 repetition and requires less inhibition. Therefore, when switching back to the L2, less residual inhibition needs to be overcome, regardless of whether the run was short or long. Alternatively, the L2 requires more enhancement during its repetition, hence more enhancement needs to be overcome when switching to L1.

A different view on the dynamics of language control has been proposed by Kleinman and Gollan (2018), who argued that inhibition accumulates over time. Using the same picture-naming paradigm, they tracked how naming RTs of the target picture changed as a function of the number of unrelated pictures having been named in the alternative language. Crucially, they considered the increase of RTs *within a mixed-language block* as an index of inhibition, rather than the RTs *within a consecutive same-language run*, as investigated in Zheng et al. (2018). Their results showed that the more unrelated pictures bilinguals had previously named in the nontarget language, the slower they became in naming pictures in the target language. This global inhibition effect was only found in the L1, but not in the L2. The authors argued that every retrieval in the nondominant L2 hinders subsequent retrieval in the dominant L1, but not vice versa. Interestingly, the run-length effect observed by Zheng et al. was also replicated by Kleinman and Gollan in their study, although its interaction with language was absent.

Does inhibitory control accumulate or decrease over the time course of language switching and repetition? And is it inhibition of the nontarget language, or enhancement of the target language, that drives the run-length effect? Evidence from RTs seems to be limited in this case, as there is no neutral condition to distinguish inhibition from enhancement. New insights can be gathered with the help of EEG, where inhibitory control in language switching is often associated with an N2 effect.

The N2 in event-related potentials (ERPs) is a negative-going component peaking around 200 to 350 ms after stimulus onset. It is commonly associated with response inhibition, such as withholding the button press in a go/no-go task (Falkenstein, Hoormann, & Hohnsbein, 1999; Jodo & Kayama, 1992). With respect to language switching, a larger N2 has been observed for switch trials compared to repeat trials, with the switch-cost effect only presented in the L2 (Jackson, Swainson, Cunnington, & Jackson, 2001). This N2 switch effect has a fronto-central scalp distribution which is similar to the no-go N2 (but see Christoffels et al., 2007, for a report where a larger N2 was found on repeat trials than on switch trials, particularly in L1). Jackson et al. (2001) argued that the N2 effect on switching reflected inhibition of the competing nontarget lexicon; greater inhibition of L1 is required when switching to L2 compared to switching to L1. Interestingly, in the same study, a larger switch cost in RT was observed in the L1 rather than in the L2. To explain the difference between the RT and the ERP results, the authors argued that “the frontal N2 reflects processes that are in operation to bring about switching whereas the RT data reflect the net result of having switched” (Jackson et al., 2001, p. 177). A later study replicated the N2 effect for switch costs (to L2) in bilingual picture-naming (Verhoef, Roelofs, & Chwilla, 2010), but the reported N2 had a more posterior rather than anterior scalp distribution (see also Folstein & Van Petten, 2008, for a review of the dissociation between the anterior and posterior N2 in the nonlinguistic literature). This posterior N2 effect was interpreted as to reflect the disengagement of the nontarget language: Switching to the L2 requires disengagement of the stronger L1 (therefore a larger N2 effect), which is not the case for switching to the L1 (therefore a smaller or no N2 effect). A similar posterior N2 effect has been reported in monolingual task switching as well (Sikora, Roelofs, & Hermans, 2016). In that study, speakers were asked to switch between describing black-and-white pictures with short phrases (“the fork”) and describing colored pictures with long phrases (“the green fork”). Because the short phrases need to be inhibited during the production of long phrases, it was more difficult to overcome such inhibition when participants had to switch back to the short phrases compared to the reverse situation. Therefore, a larger posterior N2 effect was observed during switches to short compared to long phrases. We will get back to the similarities and differences between these studies in the discussion.

The interplay between the anterior and posterior N2s may shed light on the question how inhibitory control develops during repeated use of the same language and how such inhibition is overcome during language switching. To this end, we employed the bilingual picture-naming paradigm used in Zheng et al. (2018) and measured bilingual speakers’ EEG during naming. To examine whether inhibitory control accumulates or decreases during language repetition, we compared responses on early vs. late ordinal positions within a same-language run.^1^ If the target language gets increasingly activated throughout repetition, we should expect faster responses on late than early ordinal positions. Furthermore, if inhibitory control decreases due to bottom-up activation of the target language, then a decrease in the N2 amplitude, as an index of inhibition, should be observed in late compared to early ordinal positions as well. We expected this N2 effect, if observed, to have an anterior scalp distribution, which is associated with response inhibition (Falkenstein et al., 1999) or language inhibition (Jackson et al., 2001).

Besides the investigation of repeat trials, we also looked at switch trials to answer the question whether the run-length effect (i.e., switching is more costly following a short compared to a long same-language run) is due to overcoming inhibition at the switch. To this end, we compared the N2 at switch trials following short vs. long same-language runs. If the process of overcoming inhibition dominates during switching, we would expect to replicate Zheng et al. (2018) in the behavioral results by finding a larger switch cost in RTs when switching after a short run compared to a long run. Furthermore, a larger N2 should be observed at switches following a short than a long run. The scalp distribution of the N2 switch effect may be either more anterior, reflecting inhibition (Jackson et al., 2001), or more posterior, reflecting disengagement or overcoming inhibition (Sikora et al., 2016; Verhoef et al., 2010).

## Method

### Participants

Thirty participants took part in the study for course credit or vouchers. All of them were native Dutch speakers, raised monolingually, who spoke English as their most proficient nonnative language. All participants were right-handed and had normal or corrected-to-normal vision. Participants were recruited online using the Radboud research participation system and received study credits or vouchers for compensation. The study was conducted in accordance with the Declaration of Helsinki, was approved by the local ethics committee (Faculty Ethics Committee, Radboud University, ECSW2015-2311-349), and all subjects provided written informed consent.

Four participants’ data were excluded from the EEG analysis due to excessive artifacts, and one additional participant was excluded due to a technical problem during recording. To be consistent, we also excluded their data from the behavioral analysis. This resulted in a final set of 25 participants (seven males).

Table 1 summarizes the language background of the 25 participants as assessed by a questionnaire, and their English vocabulary size measured by the LexTALE test (Lemhöfer & Broersma, 2012).

**Table 1:**
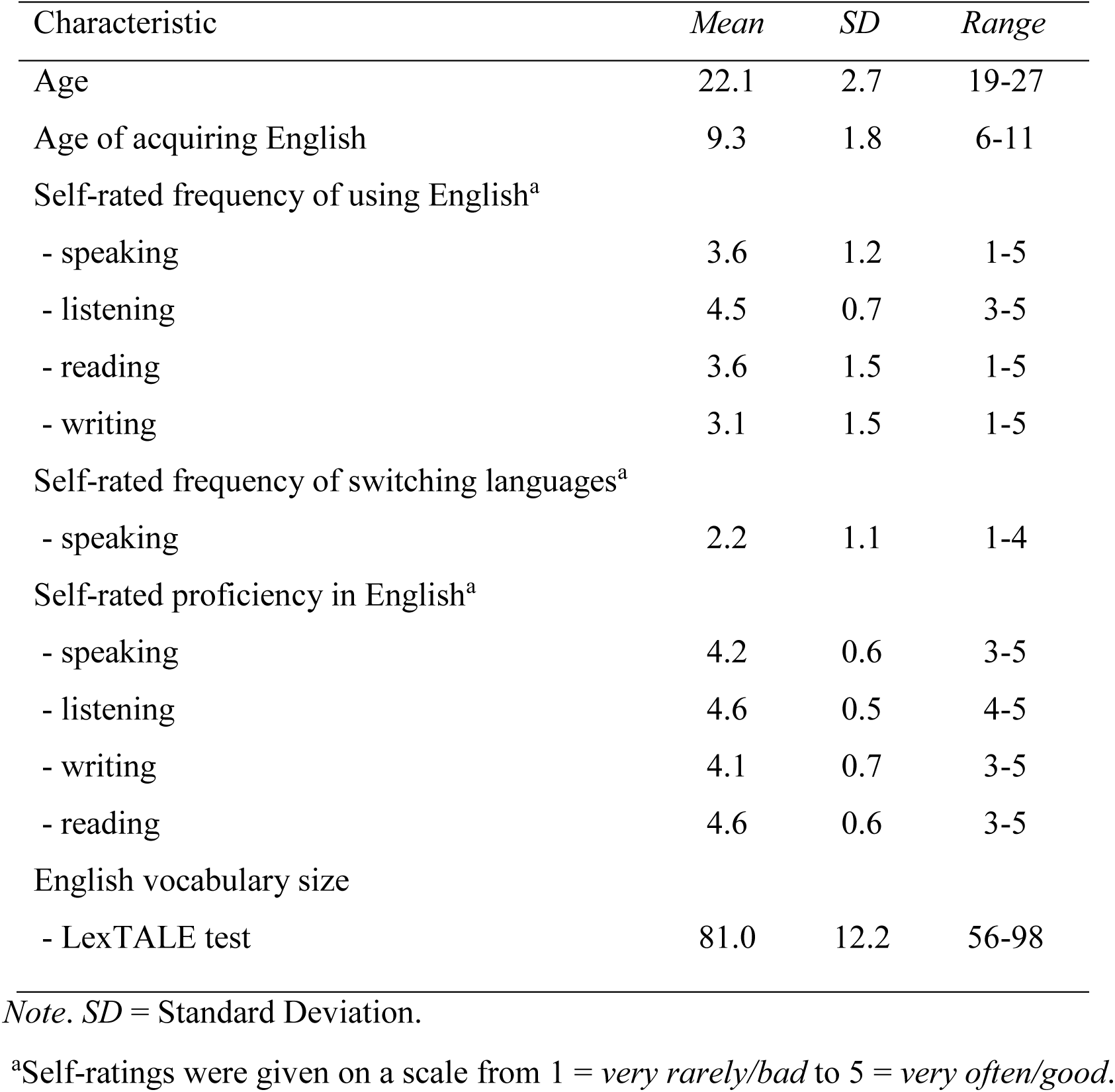
Participants’ language background and English proficiency

| Characteristic | *Mean* | *SD* | *Range* |
| --- | --- | --- | --- |
| Age | 22.1 | 2.7 | 19-27 |
| Age of acquiring English | 9.3 | 1.8 | 6-11 |
| Self-rated frequency of using English<sup>a</sup> |  |  |  |
| - speaking | 3.6 | 1.2 | 1-5 |
| - listening | 4.5 | 0.7 | 3-5 |
| - reading | 3.6 | 1.5 | 1-5 |
| - writing | 3.1 | 1.5 | 1-5 |
| Self-rated frequency of switching languages<sup>a</sup> |  |  |  |
| - speaking | 2.2 | 1.1 | 1-4 |
| Self-rated proficiency in English<sup>a</sup> |  |  |  |
| - speaking | 4.2 | 0.6 | 3-5 |
| - listening | 4.6 | 0.5 | 4-5 |
| - writing | 4.1 | 0.7 | 3-5 |
| - reading | 4.6 | 0.6 | 3-5 |
| English vocabulary size |  |  |  |
| - LexTALE test | 81.0 | 12.2 | 56-98 |

*Note.* *SD* = Standard Deviation.
<sup>a</sup>Self-ratings were given on a scale from 1 = *very rarely/bad* to 5 = *very often/good*.

### Materials

Forty black-and-white line drawings, which represented 40 translation pairs of Dutch–English noncognate words (e.g., the Dutch word “boom” and its English translation “tree”), were used as experimental pictures (see Appendix A). All the pictures were taken from the international picture naming project (IPNP) database (Bates et al., 2003). Based on a pilot study on naming agreement, we replaced two of them with drawings sketched by the first author. Pictures were selected with high naming agreement in both Dutch and English (Bates et al., 2003; Severens, Van Lommel, Ratinckx, & Hartsuiker, 2005) and high naming frequency (CELEX database; Baayen, Piepenbrock, & Gulikers, 1995) as selection criteria. We matched all the Dutch and English picture names as closely as possible on number of syllables (*p* = .813) and phonological onset category, so that possible differences in RTs could not be explained by word length or differences in voice-key sensitivity (e.g., /f/ has a delayed voice-key onset compared to /a/). All the pictures were scaled to 300 × 300 pixels.

### Design

There were two types of trials: *switch trials*, where the response language was different from that in the previous trial, and *repeat trials*, where the response language stayed the same. On repeat trials, we compared early vs. late ordinal positions within a same-language run. In the current study, a same-language run had a maximum of six trials. Therefore, we coded trials with the ordinal position 2 and 3 within a run as early position, and trials 4, 5, 6 were coded as late position (position 1 is a switch). On switch trials, we compared naming when switching after short vs. long same-language runs. The run length could be short (i.e., two or three repeat trials before a switch) or long (i.e., five or six repeat trials). Each type of run length occurred an equal number of times. Overall, 23.75% of trials in the experiment were switch trials. A schematic diagram of the experimental paradigm can be found in Figure 1.

**Figure 1.**
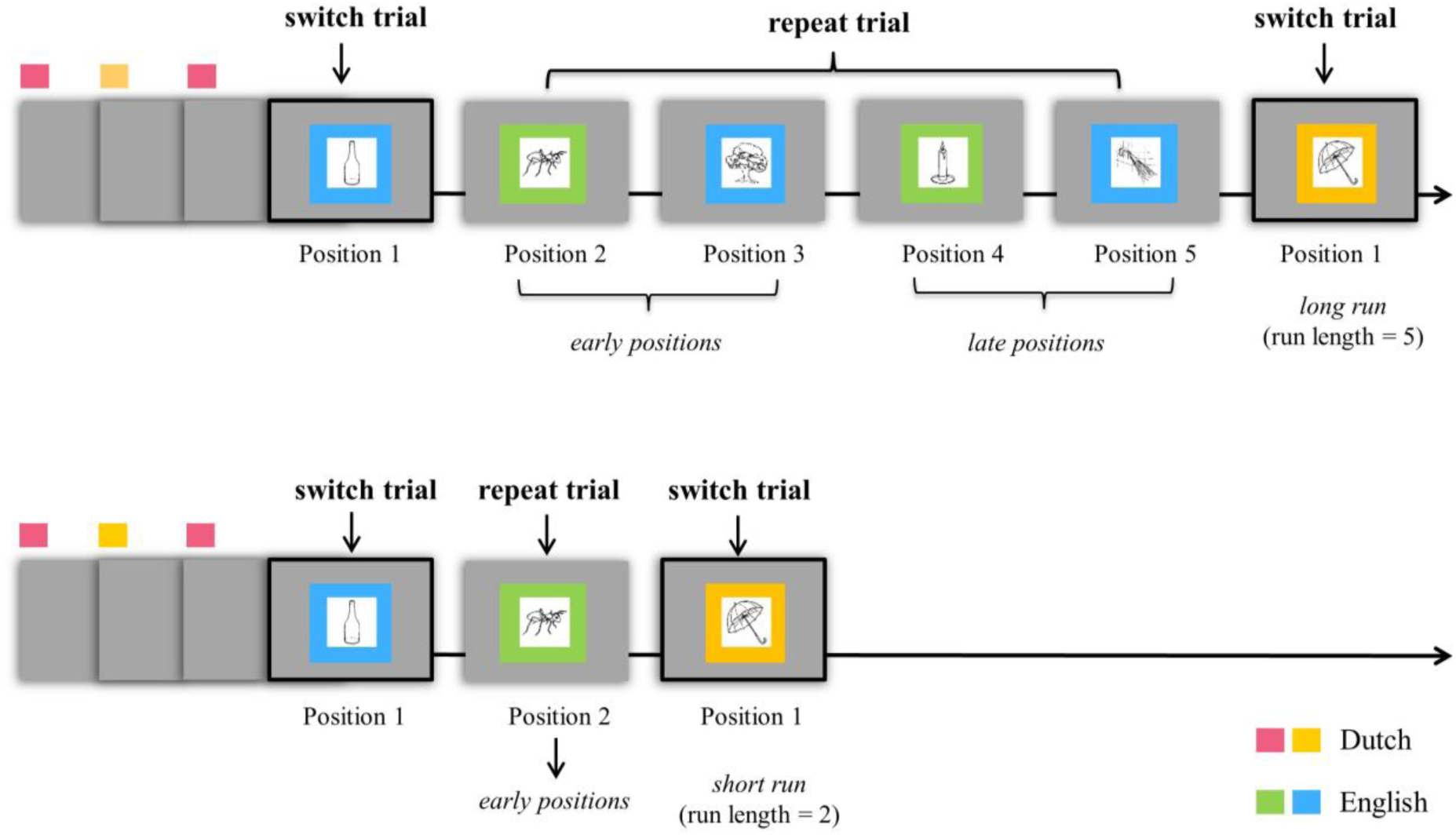
Experimental paradigm. The length of a run of same-language trials before a switch is short (run length = 2 or 3) or long (run length = 5 or 6). Trials within a same-language run are categorized as early (ordinal position = 2 or 3) or late (ordinal position = 4, 5, 6).

Each experimental list had eight blocks of 80 trials each, in total 640 trials. Each stimulus appeared twice in a block, once in Dutch and once in English. We tried to make sure that each stimulus occurred equally often on a switch trial in both languages and after all run lengths.^2^ The pseudo-randomization of repeat trials was done in each block using the program MIX (van Casteren & Davis, 2006), with the following restrictions: (1) subsequent trials were semantically and phonologically unrelated; (2) repetition of a picture was separated by at least four intervening trials. We also made sure that each item occurred at least once in each type of ordinal positions. A second list was constructed by reversing the block order of the first list.

### Procedure

Participants were seated in a sound-proof booth and the experiment was run using the software *Presentation* (Version 17.0, Neurobehavioural System Inc, Berkeley, U.S.). The background color of the computer screen (Benq XL2420Z, 24-inch screen) was set to grey, with a resolution of 1920 × 1080 pixels, at a refresh rate of 120 Hz. We first familiarized participants with all the pictures. They were asked to name each picture once in Dutch (block 1) and once in English (block 2); if they were unable to name it, they were told the correct answer and asked to remember it and name it again (block 3). Then we followed with a practice block to familiarize participants with the language cues. The cues were presented as a 100-pixel-wide frame around the picture whose color represented the response language (i.e., red and yellow for Dutch, and green and blue for English, or vice versa). Two colors were used to cue each language to avoid a confound of color switch in the stimulus and language switch in the required response (Mayr & Kliegl, 2003). We counterbalanced the assignment of colors to the response language across participants. The practice block ran for a minimum of 40 trials and stopped when participants’ accuracy reached 90%. For the first 20 trials, participants received feedback for the correct response after each trial.

EEG was recorded during the eight experimental blocks of the main experiment. Each trial started with a fixation cross (250 ms), followed by a jittered blank screen (250-500 ms). The picture then appeared in the center of the screen together with the color cue, waiting for a response which would be registered by a voice key (Shure SM-57 microphone). After a valid response or no response within a time limit (2000 ms), the stimulus stayed on the screen for another 550 ms. The next trial began after another jittered blank screen (250-500 ms). We instructed the participants to name the pictures as quickly as possible in the language indicated by the cue, and also not to correct themselves when they said something wrong. All the instructions were given in Dutch.

After the main experimental part with EEG measurement, participants completed the LexTALE vocabulary test in English and a language background questionnaire. The entire session took approximately 2 hrs.

### EEG Recording

We recorded EEG from 57 active Ag-AgCl electrodes mounted in an elastic cap, placed according to the international 10-20 system (ActiCAP 64Ch Standard-2, Brain Products). EEG signals were referenced online to the left mastoid electrode and re-referenced offline to the average of the right and left mastoid electrodes. EOG was measured horizontally with two additional electrodes placed above and below the right eye, and vertically with two electrodes placed on the left and right temples. EMG^3^ was measured with two electrodes placed next to the upper lip and the throat. EEG, EOG and EMG signals were amplified with BrainAmps DC amplifiers (500 Hz sampling, 0.016 – 125 Hz band-pass). Impedances for EEG electrodes were kept below 20 kΩ.

### EEG Preprocessing

We performed all EEG analyses using the Fieldtrip open source Matlab toolbox (Oostenveld, Fries, Maris, & Schoffelen, 2011) and custom analysis scripts in Matlab v.8.6.0 (R2015b, The Math Works, Inc). We first segmented the continuous EEG into epochs from 200 ms before to 2500 ms after picture onset.^4^ The data were then re-referenced and band-pass filtered with a low cut-off of 0.1 Hz and a high cut-off of 30 Hz. Trials with atypical artifacts (e.g., jumps and drifts) were rejected by visual inspection; EOG artifacts (eye blinks and saccades) were removed using independent component analysis. After that, we further segmented the data into shorter epochs from 200 ms pre- to 500 ms post-picture onset and applied another round of visual inspection to remove trials with remaining artifacts (e.g., muscle artifacts due to early articulation). Baseline correction was applied based on the average EEG activity in the 200 ms interval before picture onset. Individual EEG channels with bad signals were interpolated by a weighted average of the data from neighboring channels of the same participant. On average, we discarded 3.5% of the epochs and 1.5% of the channels. Two channels (FT7, TP7) that were interpolated in more than two participants were excluded from the group-level analyses. We averaged all the epochs for each condition and each participant. Four participants with less than 20 remaining trials in any condition were excluded from the EEG analysis (see also “Participants”).

### Statistical Analyses

Participants’ responses were categorized as errors when they used nontarget words (from either language), or when they failed to respond or respond with a repair or disfluency. Errors were excluded from the subsequent RT and ERP analyses. Naming RTs were recorded online using a voicekey and later manually corrected if necessary, using the speech analysis program Praat (Boersma & Weenink, 2016). Correctly responded trials with a RT deviating more than three standard deviations from the respective participants’ condition mean were excluded (per language and per trial type). Trials in the beginning of each block and post-error trials were also excluded. Because a language selection error on a repeat trial (e.g., saying the Dutch word “boom” instead of the English word “tree”) alters the characteristics of the run in which it occurs (e.g., turns a long run into a short run), we decided to exclude all runs with errors. This led to an exclusion of 9.9% of the data. We did not analyze the errors trials themselves due to their infrequent occurrence (3.9% on the switch trials, 1.8% on the repeat trials, before excluding all runs with errors).

The statistical analyses of the behavioral data were computed with generalized mixed-effects models using the lme4 package (Version 1.1.13, Bates, Mächler, Bolker, & Walker, 2015) in R (Version 3.4.1; R Core Team, 2017) to account for the right-skewed shape of the RT distribution without the need to transform and standardize the raw data (Lo & Andrews, 2015). We started with a full model of RTs as a function of language (L1 vs. L2), trial type (switch vs. repeat) and run length (short vs. long)/ordinal position (early vs. late), and followed up all the interactions with trial type. To further test our hypotheses on ordinal positions and run length, we also analyzed the RTs of repeat and the switch trials separately, with language and ordinal positions/run length as factors. For all the analyses, the factors language, ordinal position/run length and trial type (if applicable) were sum-coded and included as fixed effects. Participants and items were included as random effects. We ran all the models with a maximal random-effects structures, which included random intercepts and random slopes for all fixed effects and their interactions for both participants and items (Barr, Levy, Scheepers, & Tily, 2013). Only when the model with the maximal random-effects structure did not converge, we simplified it by first removing the interactions and if necessary the main effects in the random structure.

The statistical analysis of the ERP data was run using a nonparametric cluster-based permutation test (Maris & Oostenveld, 2007) using Matlab v.8.6.0 (R2015b, The Math Works, Inc). The method controls for the false alarm rate caused by multiple comparisons, i.e., when evaluating the ERP data at multiple channels and multiple time points. On the repeat trials, we compared early vs. late positions within a same-language run; on the switch trials, we compared the switch after short vs. long runs. We also compared the switch costs (i.e., difference between switch and repeat) between languages and between run lengths.

For the cluster-based permutation test, the two conditions of interest were first compared using a paired-samples *t*-test (two-tailed) at each spatiotemporal sample (i.e., per channel and time point). Then we used an alpha threshold of .05 and all samples with smaller *p*-values are selected. Afterwards, those selected samples which were spatiotemporally adjacent were grouped as clusters. For each cluster, the sum of the *t*-values of all the samples was used as the cluster-level statistic. Using the same procedure as described above, we constructed a permutation distribution by randomly partitioning the original data for 1000 times and then computing spatiotemporal clusters with their cluster-level statistic. We selected the cluster with the maximum cluster-level statistic to compare against the permutation distribution. The *p*-value of the cluster was calculated as the proportion of random partitions (out of 1000) that yielded a larger cluster-level statistic than its own statistic. A *p-*value below .05 (two-tailed) was considered to be significant.

We focused our ERP analysis on the N2 components. Following the N2 literature, statistical tests were applied to the time window of 200 ms to 350 ms post stimulus onset. Given the two possible topographies of the N2 components (i.e., anterior N2 and posterior N2), we applied our analysis to all available electrodes.

## Results

### Behavioral Results

#### Overall analysis

Figure 2 shows the violin plots for the RTs on the repeat trials (top panel) and on the switch trials (bottom panel).

**Figure 2.**
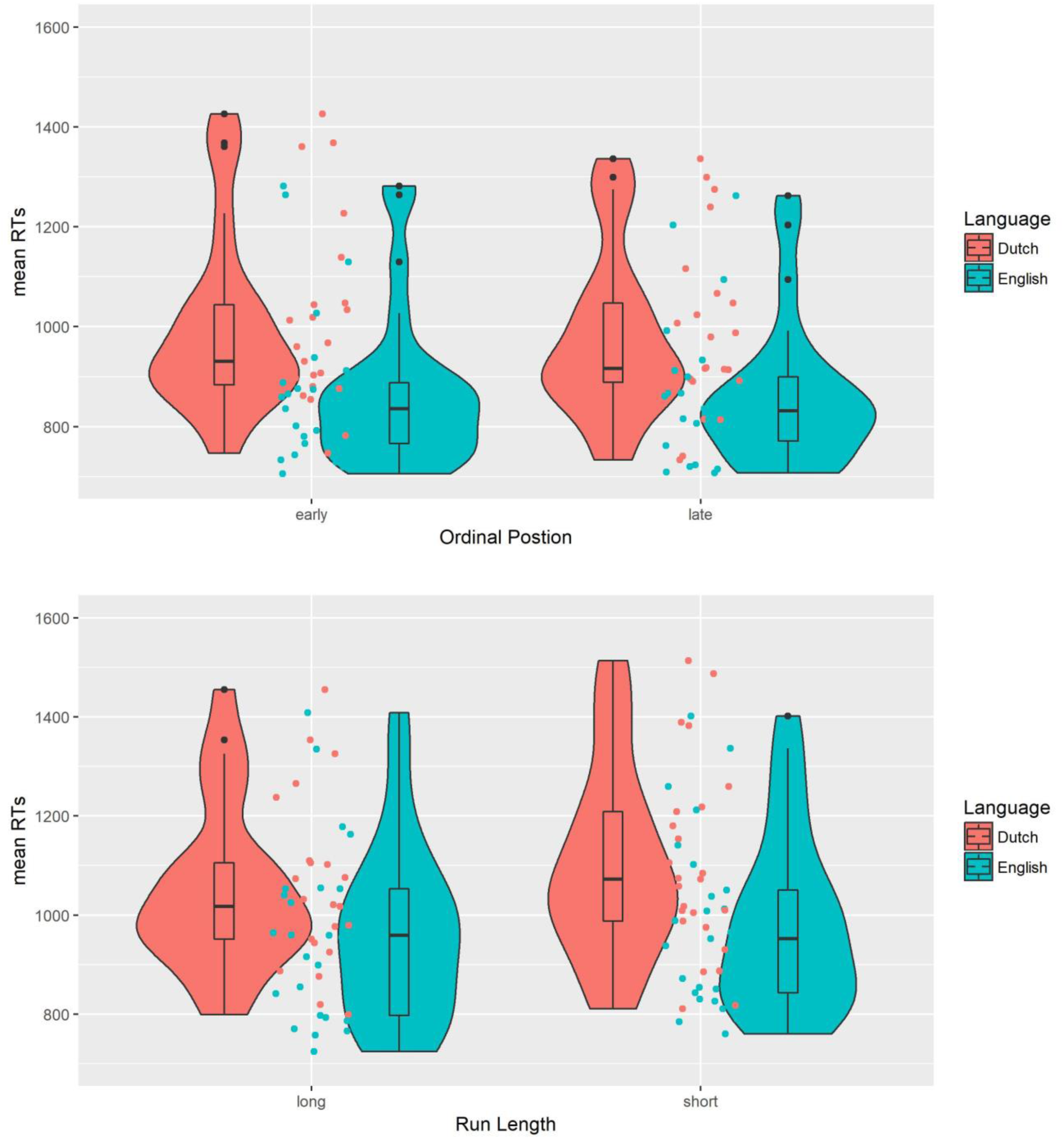
Violin plots with individual data distributions of mean RTs (in ms) for repeat trials (top panel) and switch trials (bottom panel), grouped by language (Dutch vs. English) and ordinal position (early vs. late, top panel) or run length (long vs. short, bottom panel). The outer shapes represent the distribution of individual data, the thick horizontal line inside the box indicates the median, and the bottom and top of the box indicate the first and third quartiles of each condition.

We started with a full model of RT as a function of language (L1 vs. L2), trial type (switch vs. repeat), and run length/ordinal position. For the run length/ordinal position analysis, early vs. late position of repeat trials were used as a baseline for short vs. long runs of switch trials, respectively. Table 2 presents all the statistics from the GLMEM used for this analysis.

**Table 2.**
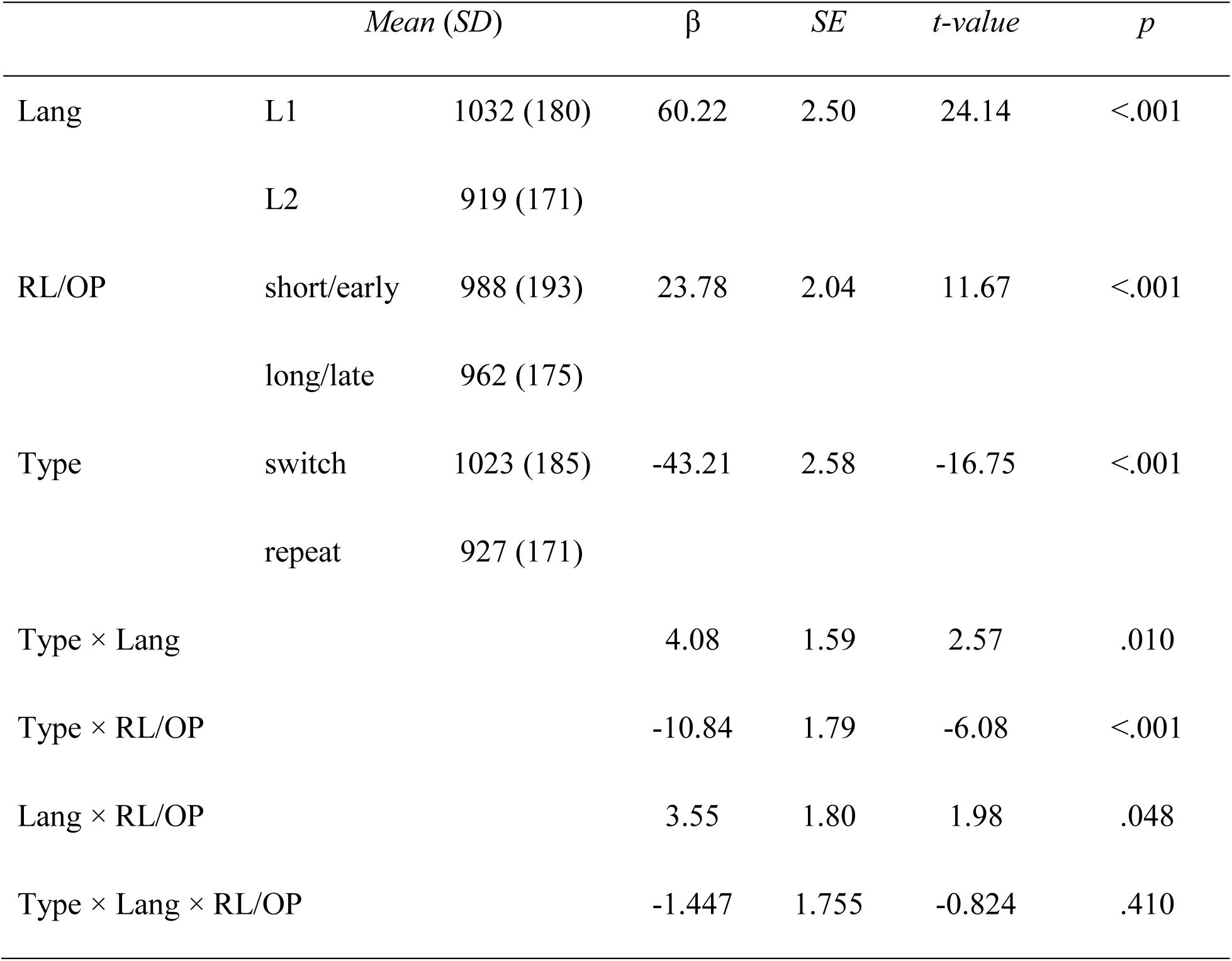
Statistics from the GLMEM for the reaction time (RT, ms) as a function of language (Lang), trial type (Type), and run length/ordinal position (RL/OP)

Bilingual speakers were slower to respond in L1 than in L2, and on switch than on repeat trials. There was also a main effect of run length/ordinal position: Speakers were faster in late positions/when switching after a long run compared to early positions/when switching after a short run.

##### Switch costs

We observed an interaction between trial type and language. Follow-up analyses showed that the switch cost (i.e., the difference between trial types) was smaller in L1 (*diff* = 87 ms; β = −41.68, *SE* = 5.06, *t* = −8.23, *p* < .001) than in L2 (*diff* = 105 ms; β = −55.08, *SE* = 4.36, *t* = −12.62, *p* < .001). Another interaction was found between trial type and run length/ordinal position: the switch cost was larger for a short run (*diff* = 107 ms; β = −56.80, *SE* = 5.76, *t* = −9.86, *p* < .001) than a long run (*diff* = 85 ms; β = −43.98, *SE* = 4.42, *t* = −9.96, *p* < .001).

To test our hypotheses on ordinal positions and run length, we further analyzed RTs separately for repeat trials and switch trials.

#### Analysis of repeat trials

On the repeat trials, we analyzed how naming RTs differed on early vs. late ordinal positions within a same-language run, and how it interacted with language. The statistics from the GLMEMs for the RTs on the repeat trials are presented in Table 3 (top panel).

**Table 3.**
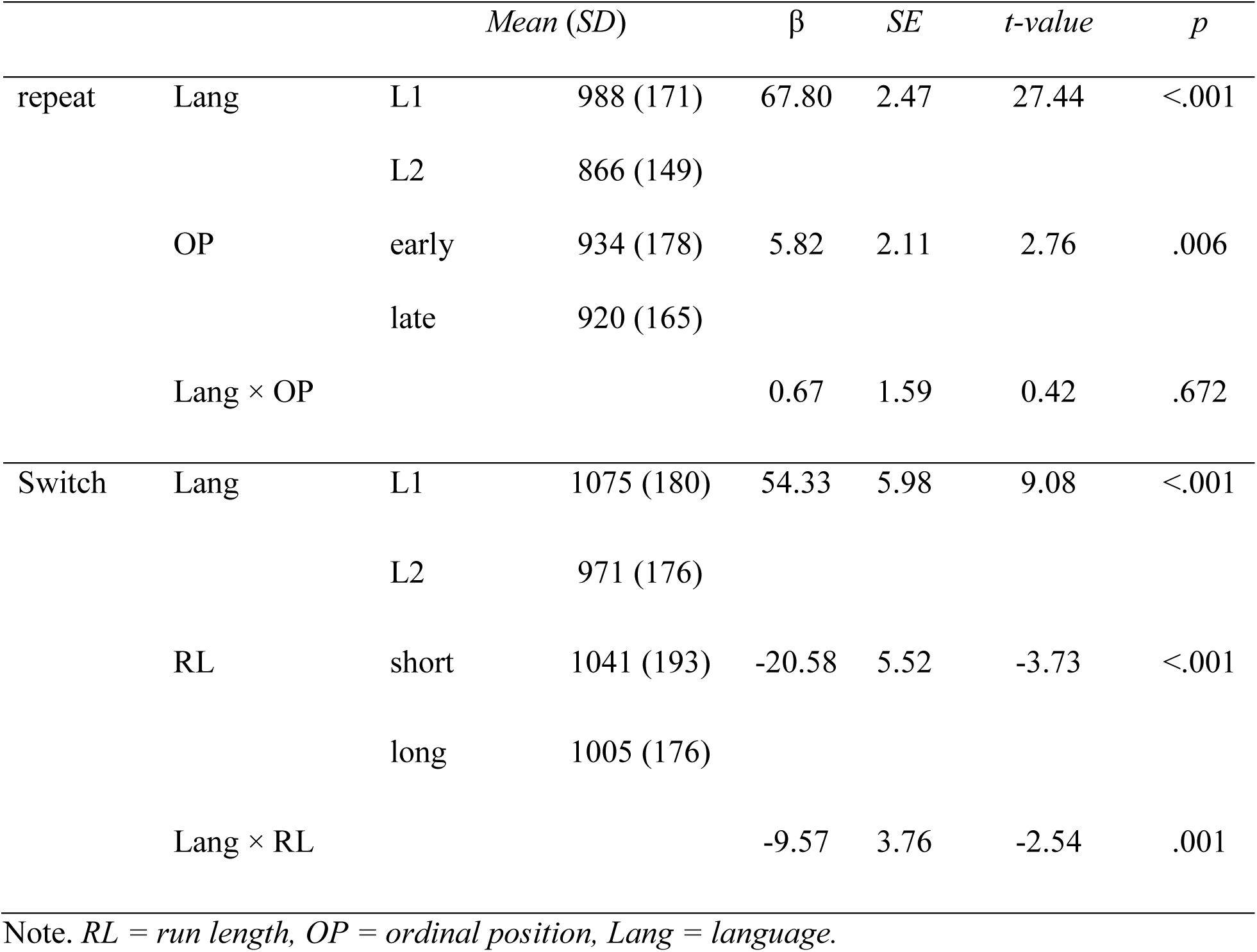
Statistics from the GLMEMs for the reaction time (RT, ms) on repeat and switch trials, respectively

| | | | Mean (SD) | $\beta$ | SE | t-value | p |
| --- | --- | --- | --- | --- | --- | --- | --- |
| repeat | Lang | L1 | 988 (171) | 67.80 | 2.47 | 27.44 | <.001 |
|  |  | L2 | 866 (149) |  |  |  |  |
|  | OP | early | 934 (178) | 5.82 | 2.11 | 2.76 | .006 |
|  |  | late | 920 (165) |  |  |  |  |
| | Lang × OP | | | 0.67 | 1.59 | 0.42 | .672 |
| Switch | Lang | L1 | 1075 (180) | 54.33 | 5.98 | 9.08 | <.001 |
|  |  | L2 | 971 (176) |  |  |  |  |
|  | RL | short | 1041 (193) | -20.58 | 5.52 | -3.73 | <.001 |
|  |  | long | 1005 (176) |  |  |  |  |
| | Lang × RL | | | -9.57 | 3.76 | -2.54 | .001 |

Note. RL = run length, OP = ordinal position, Lang = language.

On repeat trials, speakers were slower in the L1 than the L2, and in early than late positions. There was no interaction between language and ordinal positions.

#### Analysis of switch trials

On switch trials, we analyzed how naming RTs differed in switches after short vs. long same-language runs, and how this interacted with language. Table 3 (bottom panel) gives the statistics from the GLMEMs for the RTs on the switch trials.

Speakers were slower to switch after a short run than a long run, and when switching to the L1, Dutch, than to the L2, English. There was a significant interaction between language and run length: The run-length effect was only present in the L1 (*M*_L1short_ = 1101 ms, *SD*_L1short_ = 193 ms; *M*_L1long_ = 1049 ms, *SD*_L1long_ = 167 ms; β = −28.27, *SE* = 8.28, *t* = −3.42, *p* < .001), but not in the L2 (*M*_L2short_ = 981 ms, *SD*_L2short_ = 177 ms; *M*_L2long_ = 961 ms, *SD*_L2long_ = 177 ms; β = −10.94, *SE* = 8.79, *t* = −1.24, *p* = .213).

#### Summary

Speakers were slower in the L1 than in the L2 – replicating the reversed dominance effect – and when switching compared to repetition of language. Interestingly, switch cost was larger in L1 than in L2, which seems to be contradictory to previous literature (e.g., Meuter & Allport, 1999). We will address this in the Discussion. On repeat trials, speakers were faster in later than early ordinal positions, suggesting bottom-up activation of the target language. On switch trials, speakers were faster to switch after a long run than a short run. The run-length effect was only present in the L1, not in the L2, replicating Zheng et al. (2018).

### ERP Results

#### Analysis of repeat trials

Figure 3 shows the averaged ERPs and topographies for early vs. late ordinal positions within a same-language run, in three representative midline electrodes: Fz (anterior), Cz (central), and Pz (posterior).

**Figure 3.**
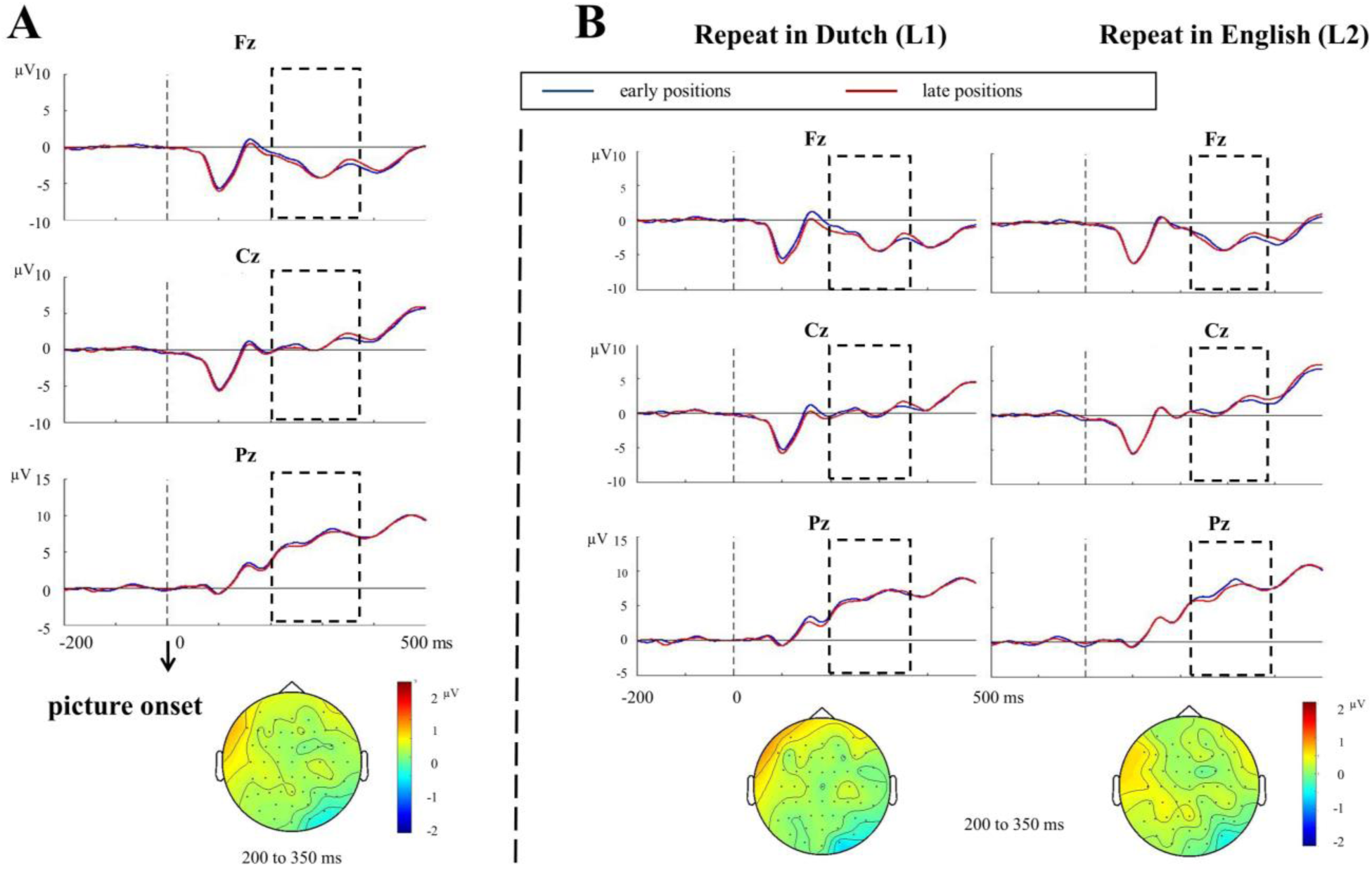
(A) Stimulus-locked ERPs and topographies for early vs. late ordinal positions, averaged across the two languages. (B) Stimulus-locked ERPs and topographies for early vs. late ordinal positions when repeating in L1 (Dutch) and repeating in L2 (English). The time window used for testing the N2 effect (200 to 350 ms) is marked by a dotted frame. Topographies of the difference between the two conditions within the time window for testing are presented for each contrast.

The cluster-based permutation tests showed no differential N2 in early compared to late ordinal positions (*p* = .378; Figure 3A). When further tested within each language (Figure 3B), no N2 effect was observed for early compared to late ordinal positions either in L1, Dutch (*p* = .338) or in L2, English (*p* = .308). The difference between languages was also not significant, as no clusters were detected in the permutation test.

#### Analysis of switch trials

Figure 4 shows the averaged ERPs and topographies for switch trials following short vs. long same-language runs.

**Figure 4.**
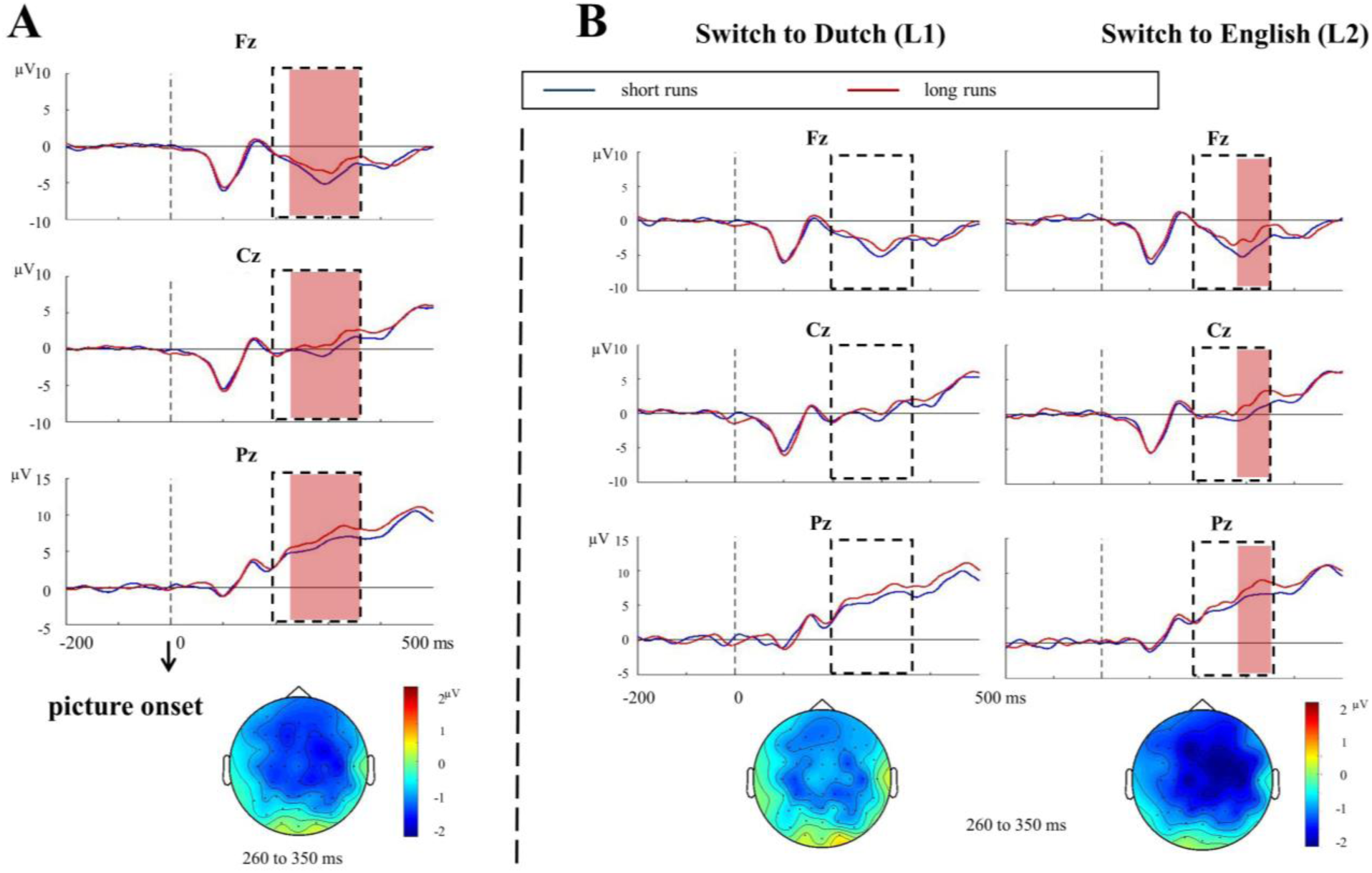
(A) Stimulus-locked ERPs and topographies for switch trials after short vs. long runs, averaged across languages. (B) Stimulus-locked ERPs and topographies for switching to L1, Dutch and to L2, English, after short vs. long runs. The time window used for testing the N2 effect (200 to 350 ms) is marked by an empty frame. When the N2 effect was significant between conditions, the time windows associated with the statistically significant effect are marked in light red. Topographies of the difference between the two conditions are presented for each contrast. We used the same time window in A and B (the window that was associated with the N2 effect in A) for depicting the topography for the sake of better comparability.

The cluster-based permutation tests revealed a significant increase in N2 amplitude on switch trials following a short compared to a long run (*p* = .010, Figure 4A). The effect was most pronounced between 260 to 350 ms post stimulus onset, with a widespread scalp distribution somewhat centered towards the fronto-central sites and right lateralized. We further compared the N2 effect between the two switching directions (Figure 4B). Results showed that the N2 run-length effect was only present when switching to the L2 (*p* = .042). The effect was most pronounced between 320 to 350 ms post stimulus onset, widely distributed over the scalp, and slightly stronger in fronto-central sites and right lateralized. In contrast, the N2 run-length effect was not observed when switching to the L1 (*p* = .178). However, this difference between languages was not significant (*p* = .360).

#### Analysis of switch cost

To better compare the current ERP results with previous literature (e.g., Verhoef et al., 2010), we contrasted the switch trials with the repeat trials (i.e., switch costs), as well as the switch costs between languages and between run lengths. Figure 5 shows the averaged ERPs and topographies for switch vs. repeat trials, and in each language, respectively.

**Figure 5.**
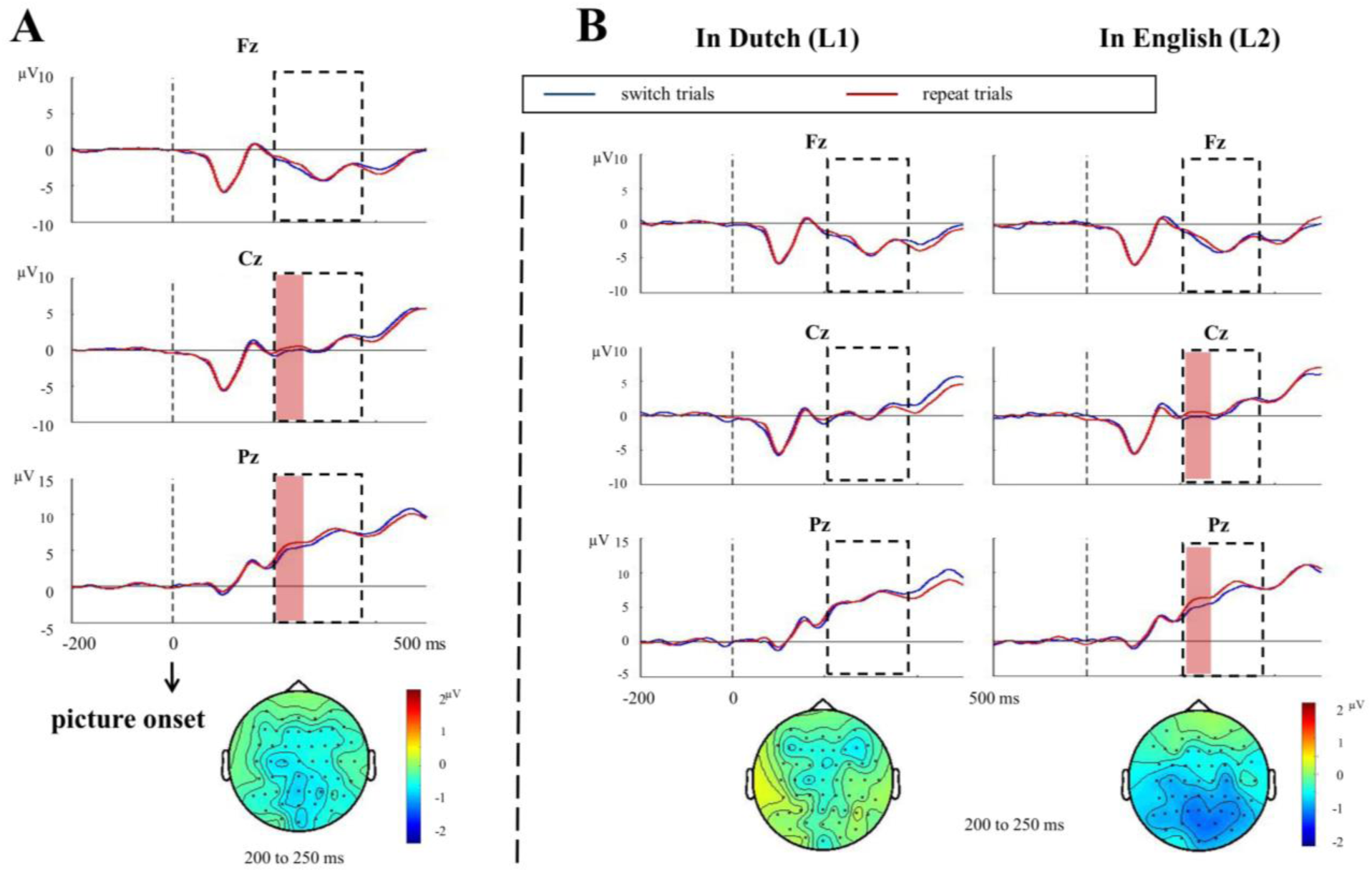
(A) Stimulus-locked ERPs and topographies for switch vs. repeat trials. (B) The same contrast (i.e., switch vs. repeat) when naming in L1, Dutch and in L2, English. The time window used for testing the N2 effect (200 to 350 ms) is marked by an empty frame. When the N2 effect was significant between conditions, the time windows and electrodes associated with the statistically significant effect are marked in light red. Topographies of the difference between the two conditions are presented for each contrast. We used the same time window in A and B (the window that was associated with the N2 effect in A) for depicting the topography for the sake of better comparability.

The cluster-based permutation test revealed a larger N2 after switch compared to repeat trials (*p* = .046), with the effect being most pronounced at centro-posterior sites, from 200 to 250 ms post stimulus onset (Figure 5A). When compared between languages, the N2 effect of switch cost was well present in L2 (*p* = .022), but not in L1 (*p* = .655). The effect in L2 was observed from 200 to 250 ms post stimulus onset, in centro-posterior sites (Figure 5B). The difference between languages, however, did not reach significance (*p* = .092).

*Switch costs following a short run*. Figure 6 shows the averaged ERPs and topographies for switch vs. repeat trials after a short run, and in each language, respectively.

**Figure 6.**
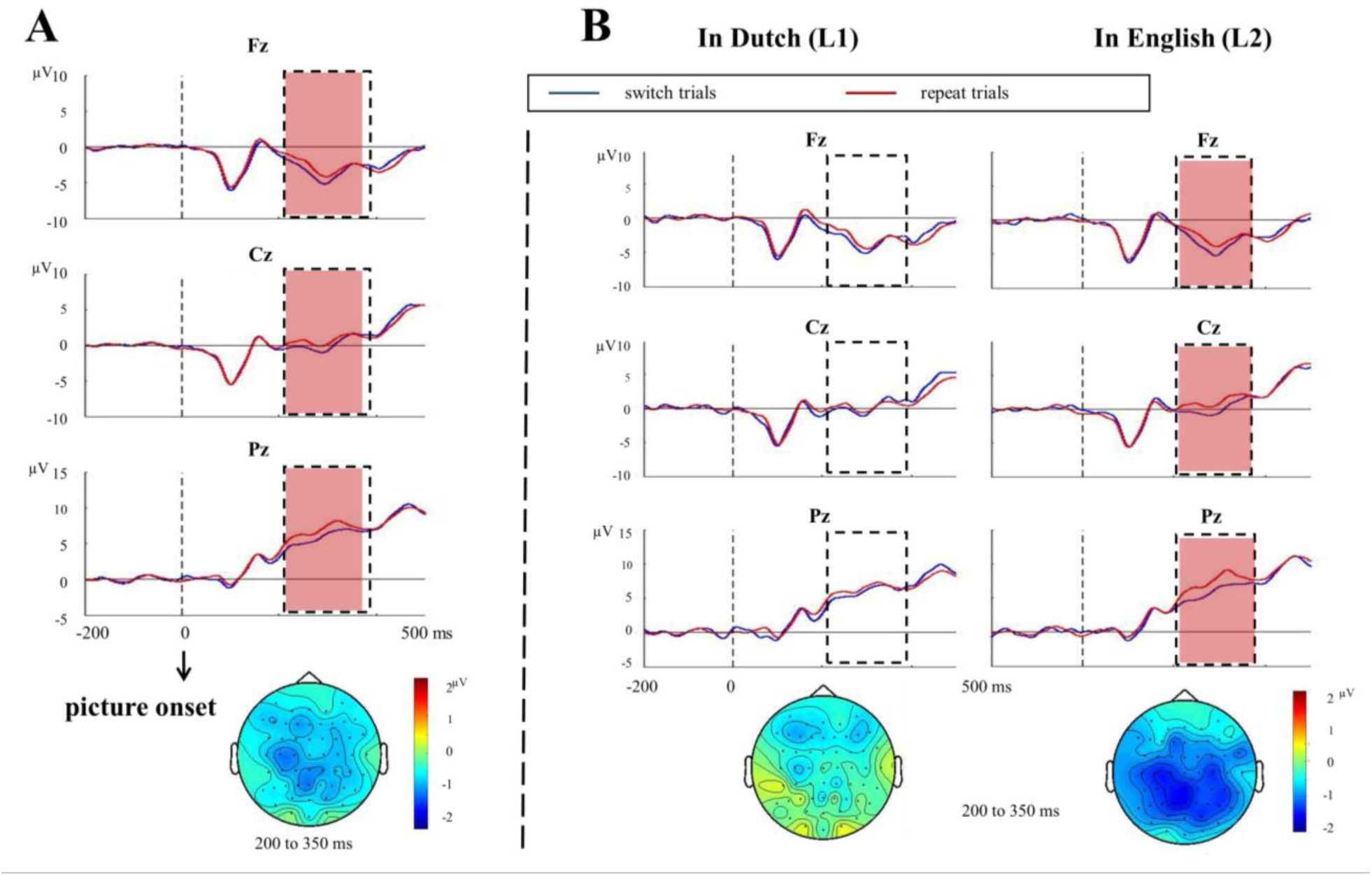
(A) Stimulus-locked ERPs and topographies for switch cost (i.e., switch vs. repeat trials) after a short run. (B) The same contrast (i.e., switch vs. repeat) when naming in L1, Dutch and in L2, English. The time window used for testing the N2 effect (200 to 350 ms) is marked by an empty frame. When the N2 effect was significant between conditions, the time windows associated with the statistically significant effect are marked in light red. Topographies of the difference between the two conditions within the time window for testing are presented for each contrast.

The cluster-based permutation tests revealed a larger N2 following the switch after a short run compared to a repeat trial of early ordinal position (*p* = .002), with the effect being most pronounced from 200 to 340 ms post stimulus onset and widely spread over the scalp (Figure 6A). When compared between languages, the switch cost N2 effect following a short run was present in the L2 (*p* = .002), but not in the L1 (*p* = .158). The effect in L2 was observed from 200 to 350 ms post stimulus onset, widely spread and more towards centro-posterior sites (Figure 6B). The difference between languages, however, was not significant (*p* = .282).

*Switch cost following a long run.* Figure 7 shows the averaged ERPs and topographies for switch vs. repeat trials after a long run, and in each language, respectively.

**Figure 7.**
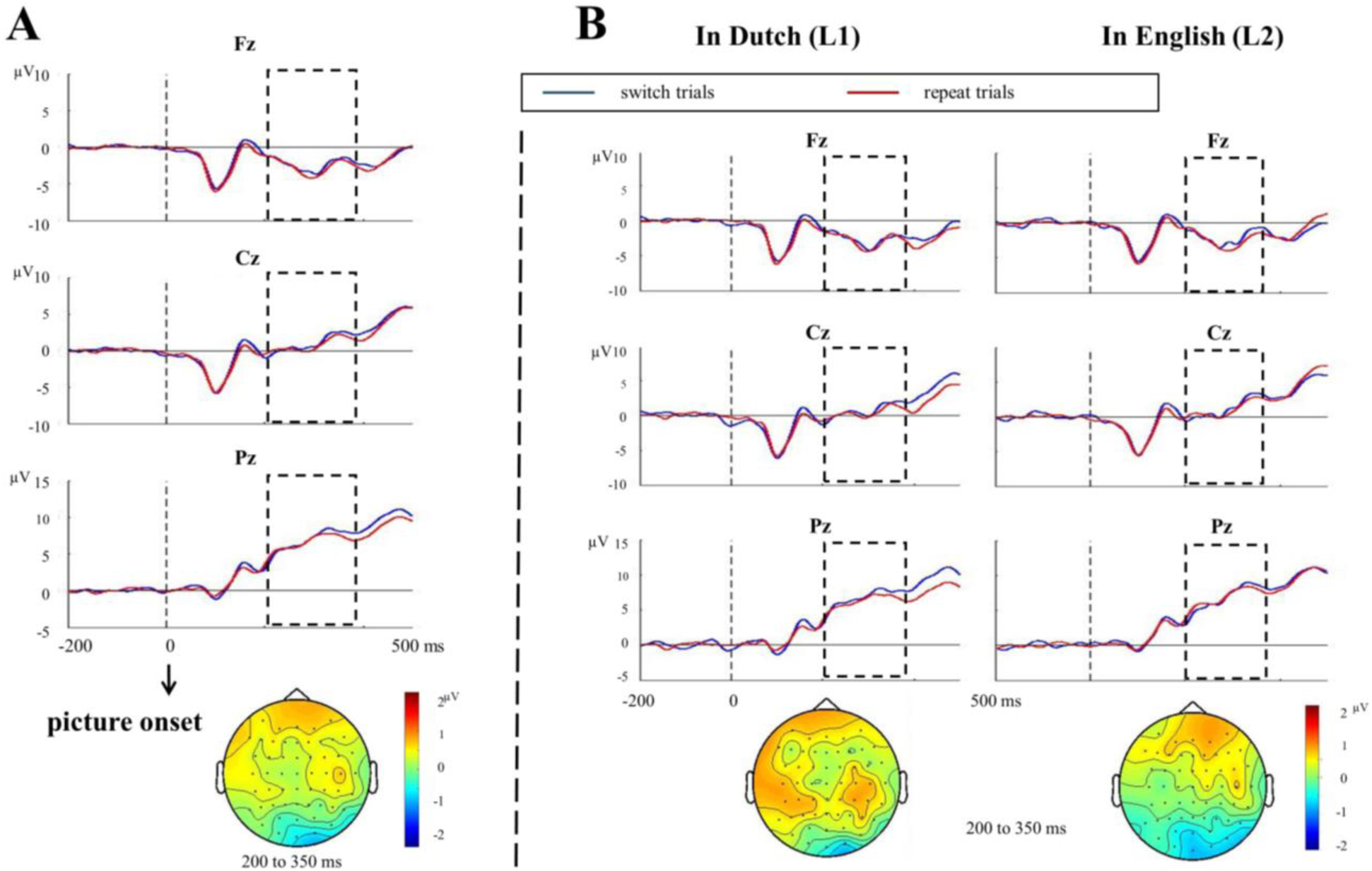
(A) Stimulus-locked ERPs and topographies for switch cost (i.e., switch vs. repeat trials) after a long run. (B) The same contrast (i.e., switch vs. repeat) when naming in L1, Dutch and in L2, English. The time window used for testing the N2 effect (200 to 350 ms) is marked by an empty frame. Topographies of the difference between the two conditions within the time window for testing are presented for each contrast.

The cluster-based permutation test showed no N2 difference between switch trials after long runs and repeat trials of late ordinal position (*p* = .320; Figure 7A). The switch cost after a long run, as reflected in the N2, was significantly different from that after a short run (*p* = .006), where a significant N2 effect was observed between trial types. When compared between languages, the switch cost N2 following a long run was neither present in the L2 (*p* = .559), nor in the L1 (*p* = .432). The difference between languages was also not significant (*p* = .482).

#### Summary

On switch trials, we observed a larger N2 after a short compared to a long run, with a wide-spread but more fronto-central scalp distribution. The N2 effect was only presented in the L2, not in the L1, although the difference was not significant between languages. In contrast, no N2 difference was observed in early vs. late ordinal positions on repeat trials. Compared to the repeat trials, the N2 was enlarged on the switch trials. The switch cost was only presented in the L2, not in the L1, with the N2 effect more pronounced at centro-posterior sites. The between-language difference was not significant, though. Moreover, the switch cost N2 was presented only after a short run, but not after a long run.

## Discussion

The current study investigated the dynamics of inhibitory control in bilingual speech production. We compared bilingual speakers’ naming RTs and ERPs between short vs. long sequences of same-language trials (i.e., run length) in a language switching task. Below, we first discuss the run length, switch cost, and reverse dominance effects in the RTs and ERPs. Next, we address the issue of whether the N2 effects reflect the application or overcoming of inhibition. Finally, the nature of top-down control, inhibition or enhancement, is discussed.

### RT results

On the switch trials, speakers were faster after a long same-language run compared to a short run. This *run-length effect* was only present in the L1, not in the L2. This directly replicated the results reported by Zheng et al. (2018; but see Meuter & Allport, 1999). The effect appears to be robust, given that the designs of our present and previous studies were not identical. For example, in Zheng et al., a combination of cognate and noncognate items was used instead of only noncognate items, as in the current study. Moreover, the research question in the previous study focused on error rates rather than RTs. To better compare the current study with the previous literature, we also calculated the switch costs, i.e., the difference between switch and repeat trials. As expected, the switch cost was larger after a short same-language run compared to a long run, indicating more effort to overcome the residual control after a short compared to a long run. On repeat trials, bilingual speakers were faster on trials in later than early ordinal positions within a same-language run, suggesting increased bottom-up activation of the target language. This also excludes the possibility that the run-length effect is due to expectation: If participants increasingly expect an upcoming switch the longer a run gets, they should get increasingly slower, rather than increasingly faster, within a same-language run.

Moreover, during the task, speakers responded more quickly when naming in their L2 rather than L1, regardless of trial type. This so-called *reversed dominance effect* has often been observed before in mixed-language contexts (Christoffels et al., 2007; Costa & Santesteban, 2004; Verhoef et al., 2010), and is probably due to the global inhibition of L1 throughout the experiment in order to facilitate L2 production, while inhibition of the generally weaker L2 is less necessary. A similar phenomenon presumably also due to greater L1 relative to L2 inhibition is called *asymmetric switch costs*, namely, switching from the L2 to the L1 takes more time than the opposite switching direction (e.g., Meuter & Allport, 1999). Surprisingly, we found a slightly larger switch cost for switches to the L2 rather than to the L1. This might be due to the presence of the reversed dominance effect. Because of the frequent language switching during the task, the difficulty of speaking L1 on a switch trial (because of the residual control) seems to be carried over to the repeat trials. In other words, it appears that under the present conditions, switch cost asymmetry evolved into a reversed dominance effect. Under a reversed language dominance, L1 turns into the “weaker” language compared to L2, and therefore, switch cost asymmetries that hinge on this dominance difference become distorted. This explanation is supported by other studies showing a reversed dominance effect in absence of asymmetric switch costs (Christoffels et al., 2007; Verhoef et al., 2010), or even with a reversed switch cost asymmetry as in our results (Declerck, Stephan, Koch, & Philipp, 2015).

Based on the behavioral results, we conclude that a target language gets more and more activated with repetition and the demand of top-down control (inhibition of the nontarget language and/or enhancement of the target language) decreases over time. However, it remains unclear whether it is inhibition or enhancement that drives the run-length effect at switches.

### ERP results

To explore the role of inhibitory control, we further investigated the N2 component in ERPs. The N2 effect in language production is usually interpreted as reflecting application of inhibition (Jackson et al., 2001) or overcoming inhibition (Sikora et al., 2016). The former usually has a frontal or central scalp distribution (anterior N2), whereas the latter has a parietal or more posterior scalp distribution (posterior N2). On repeat trials, we observed no difference in the N2 amplitude between trials in early vs. late ordinal positions. Therefore, we failed to find evidence for either accumulating or decreasing inhibition over a relative short number of language repetitions.

In contrast, on switch trials, a larger N2 was observed when switching after a short run compared to a long run, in line with our behavioral results. This N2 run-length effect had a broad scalp distribution with a more anterior rather than posterior topography. To better being able to compare our study with previous research (Jackson et al., 2001; Verhoef et al., 2010), we contrasted the ERPs between switch and repeat trials and compared the N2 switch cost between run lengths.

A small difference in N2 amplitude was observed between trial types, with more negative ERPs for switch trials compared to repeat trials, replicating earlier studies in language switching (e.g., Jackson et al., 2001; Verhoef et al., 2010; but see Christoffels et al., 2007). In line with the N2 analysis on switch trials as well as the RT results (i.e., larger switch costs after short compared to long runs), the switch cost was only present following a short run, not a long run. Different from the run-length effect in the N2 that we observed on switch trials, the topography of the “switch cost” N2 had a more posterior than frontal scalp distribution, resembling the one described in Verhoef et al. (2010) rather than Jackson et al. (2001). We will return to this point later. These results challenge the idea that inhibitory control is reactive (Green, 1998), which assumes that more activation leads to more inhibition. According to this assumption, a larger N2 should be expected at the switch following a long run, where the previous target language is more activated.

To obtain a more complete picture, we also compared the N2 as a function of trial type between languages. The switch cost N2 was only present in the L2, not in the L1, although the difference between languages was not significant. The results are in line with previous literatures (Jackson et al., 2001; Verhoef et al., 2010), but the interpretation of this asymmetry remains unclear. It can reflect more inhibition of the L1 on switches to the L2 (Jackson et al., 2001) or disengaging from the stronger L1 on switches to the L2 compared to the opposite switching direction (Verhoef et al., 2010).

### Inhibition vs. overcoming inhibition

Although the N2 effects observed in the current study usually have a widespread scalp distribution, sometimes the effect seemed to be more anterior (when contrasting switch trials following short vs. long runs), or more posterior (when contrasting the switch cost between run lengths and between languages). The anterior and posterior N2 topographies were hypothesized to reflect different control processes (i.e., inhibition vs. overcoming inhibition, respectively). However, a clear theoretical cut-off between the two is difficult. Switching is a complex process, involving multiple cognitive functions, such as shifting from the previous task to the target task, and inhibiting the nontarget task (Miyake et al., 2000). Thus, the N2 switch effect could be a combination of inhibition (anterior N2) and overcoming inhibition (posterior N2).

It is worth noting that the two studies in which a posterior N2 in language production has been reported did not use the same experimental paradigm (Sikora et al., 2016; Verhoef et al., 2010). In Sikora et al. (2016), a short phrase (e.g., “the fork”) needed to be inhibited while producing the long phrase (e.g., “the green fork”). Therefore, when switching back to the short phrase, more inhibition needed to be overcome (larger N2 effect) as compared to switching to the long phrase. Using the same logic, when switching to the L1, more residual inhibition needs to be overcome because the L1 is more strongly inhibited during L2 production. As a consequence, a larger N2 should be expected in switches to L1 than vice versa, which is neither the case in the current findings nor in previous studies (e.g., Jackson et al., 2001; Verhoef et al., 2010).

Actually, it is unclear in all these accounts what needs to be inhibited, to be overcome, or to be disengaged from. Previous literature diverges on this issue. Meuter and Allport (1999) speak about “disengagement” from the preceding language set (e.g., inhibiting the L1 and enhancing the L2 while speaking in the weaker L2) instead of disengaging from the previous target language (as proposed in Verhoef et al., 2010), which is very similar to the “overcoming inhibition” story (Sikora et al., 2016). In that scenario, the disengagement effort was reflected in the asymmetric switch costs (i.e., actively disengaging from the task set of preceding L2 repeat trials is more difficult than preceding L1 repeat trials). If such a “disengagement” effort is reflected by the N2 amplitude as well, then one should expect a larger N2 effect in switching to L1 than to L2. Again, this is opposite to what has been found in Jackson et al. (2001), Verhoef et al. (2010), and in the current study.

Another interesting observation in the current study is that the N2 run-length effect was present only in the L2. In contrast, the RT run-length effect was present in the L1 rather than L2. This might be due to a negative relationship between the amount of inhibition, as reflected by the N2 amplitude, and the RT difference: A larger N2 at the switch to the L2 after a short L1 run suggests more inhibition compared to a switch after a long run. The successful application of inhibition causes a smaller increase of the RT compared to long runs. The opposite holds for switching to L1 after short vs. long runs: less inhibition (i.e., smaller N2 effect) and therefore a larger RT difference between short and long runs. Similar results have been reported in a study by Shao et al. (Shao, Roelofs, Acheson, & Meyer, 2014). Monolingual participants were asked to name pictures with high vs. low naming agreement. During production, the alternative response candidates need to be inhibited (e.g., “sofa” for “couch”). They found that participants with successful recruitment of inhibition (as measured by a delta plot of RTs) showed a larger N2 effect as well as a smaller RT difference between conditions with high vs. low naming agreement.

### Top-down control in language repetition: inhibition or enhancement?

The question of inhibition vs. overcoming inhibition aside, it also remains unclear whether it is inhibition or enhancement that drives the run-length effect. Given that there was no ERP evidence for either decreasing or accumulating inhibition for the repeat trials, the “residual control” to be overcome on the switch trials is more likely to be the residual effect of enhancement rather than inhibition. Alternatively, it is also possible that the difference in inhibitory control between early vs. late positions in a same-language run is not large enough to be visible in the ERPs, but is large enough to make a difference at the switch where more top-down control is required, as reflected in the difference in effect between short vs. long runs.

In the study by Kleinman and colleagues (2018), naming RTs of the target picture increased within a block as a function of the number of unrelated pictures that had been named in the alternative language. This evidence for accumulating inhibition over time seems to contradict the run-length effect in the current study and in Zheng (2018) at first glance. However, such contradiction may not be too surprising. It is possible that within a same-language run, the inhibition of the nontarget language decreases due to the bottom-up activation of the target language. However, every time a switch to the alternative language occurs, the inhibition of the nontarget language needs to be brought back to a higher level. As a consequence, over the course of an entire language-mixing block, the overall inhibition of the nontarget language accumulates. Therefore, the accumulative inhibition is not (merely) caused by the use of the target language, but rather by the frequent switching between languages. Future studies can investigate the effect of switching frequency on inhibition to further investigate the dynamics of language switching and repetition.

### Summary

The current study explored the dynamics of inhibitory control during bilingual speech production by examining RTs and ERPs. We replicated the behavioral RT results as reported in Zheng et al. (2018). The results suggest that top-down control (inhibition of the nontarget language and/or enhancement of the target language) is highest at a switch. With repeated use of the same language, the target language receives more and more bottom-up activation and RT decreases. As a consequence, top-down control gets reduced over time and becomes easier to be overcome, reflected in faster switching following a long than a short run. Correspondingly, we found a larger N2 effect following short same-language runs compared to long runs, indicating more control effort in the former case. In contrast, no difference in N2 was observed within a same-language run. Our ERP results suggest that bilingual speakers adjust the level of inhibitory control during switching but not repetition of language.

## Appendix A: Stimuli

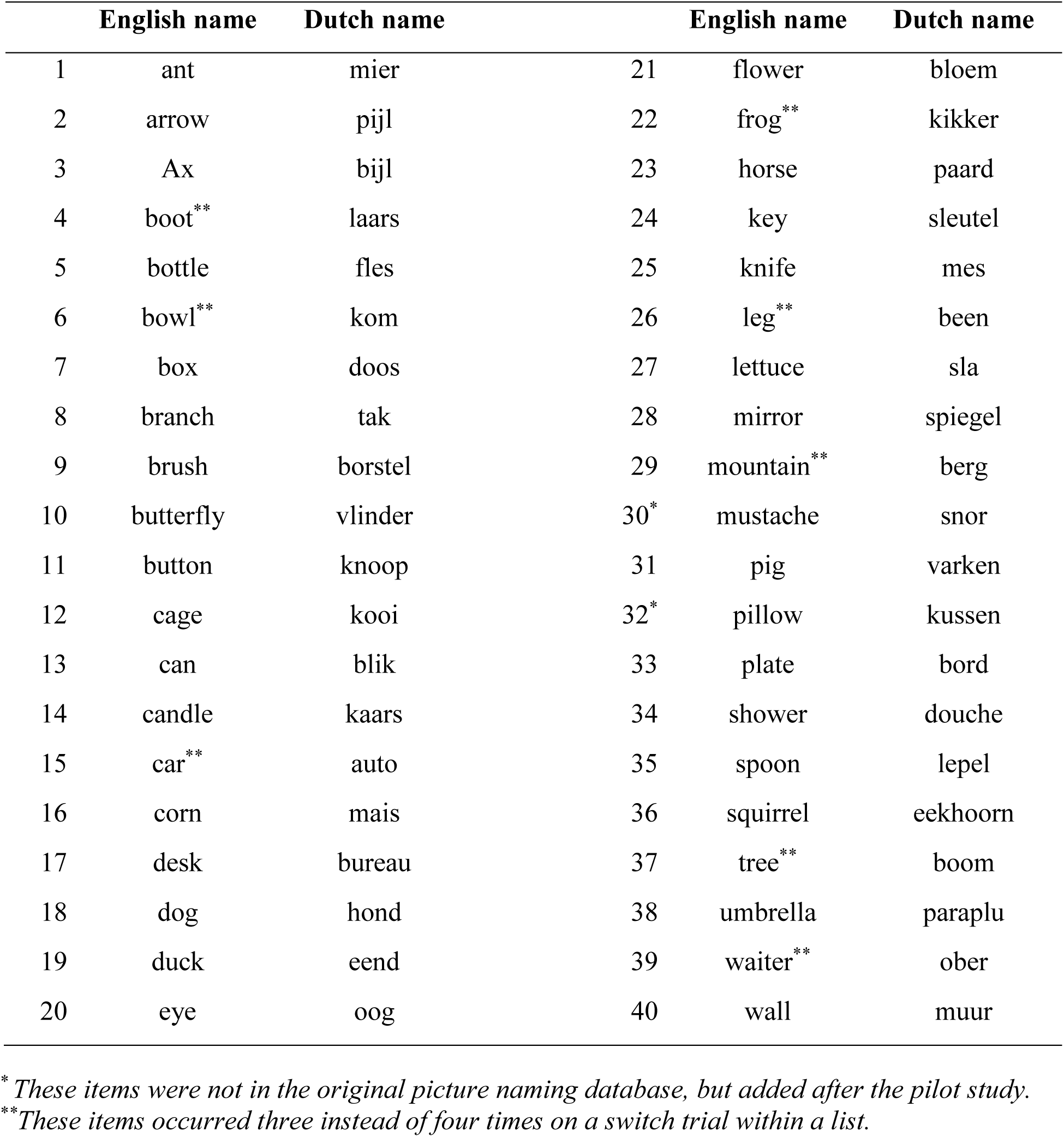

## Appendix B: Linear Mixed Effect Models

### Overall analysis

glmer.RT.all

= glmer(RT∼ TrialType*Language*RL.OP + (1+ TrialType+Language+RL.OP|pNumber) + (1+ TrialType+Language+RL.OP|PicNam), data=mydata.4RT.all, family = Gamma(link = “identity”), control=glmerControl(optimizer = ‘bobyqa’))

\# full model fails to converge

### Switch cost

\# Switch cost in L1

glmer.RT.all.swista.L1

= glmer(RT∼ TrialType + (1+ TrialType|pNumber) + (1+ TrialType|PicNam),data=mydata.4RT.all[mydata.4RT.all$Language == ‘Dutch’,], family = Gamma(link = “identity”), control=glmerControl(optimizer = ‘bobyqa’))

\# Switch cost in L2

glmer.RT.all.swista.L2

= glmer(RT∼ TrialType + (1+ TrialType|pNumber) + (1+ TrialType|PicNam),data=mydata.4RT.all[mydata.4RT.all$Language == ‘English’,], family = Gamma(link = “identity”), control=glmerControl(optimizer = ‘bobyqa’))

\# Switch cost after a short run

glmer.RT.all.swista.short

= glmer(RT∼ TrialType + (1+ TrialType|pNumber) + (1+ TrialType|PicNam),data=mydata.4RT.all[mydata.4RT.all$RL.OP == ‘short’,], family = Gamma(link = “identity”), control=glmerControl(optimizer = ‘bobyqa’))

\# Switch cost after a long run

glmer.RT.all.swista.long

= glmer(RT∼ TrialType + (1+ TrialType|pNumber) + (1|PicNam), data=mydata.4RT.all[mydata.4RT.all$RL.OP == ‘long’,], family = Gamma(link = “identity”), control=glmerControl(optimizer = ‘bobyqa’))

\# full model fails to converge

### Repeat trials

glmer.RT.repeat

= glmer(RT ∼ Language*OP + (1+Language+OP|pNumber) + (1+Language|PicNam), data=mydata.4RT.repeat, family = Gamma(link = “identity”), control=glmerControl(optimizer = ‘bobyqa’))

\# full model fails to converge

### Switch trials

glmer.RT.switch

= glmer(RT ∼ Language*RL + (1+Language*RL|pNumber) + (1+Language+RL|PicNam), data=mydata.4RT.switch, family = Gamma(link = “identity”), control=glmerControl(optimizer = ‘bobyqa’))

\# full model fails to converge

\# Interaction

glm.RT.switch.L1 =

glmer(RT∼RL + (1+RL|pNumber) + (1+RL|PicNam), data=mydata.4RT.switch.[mydata.4RT.switch$Language==“Dutch”,], family = Gamma(link = “identity”), control=glmerControl(optimizer = ‘bobyqa’))

glm.RT.switch.L2 =

glmer(RT ∼ RL + (1+RL|pNumber) + (1+RL|PicNam), data=mydata.4RT.switch[mydata.4RT.switch$Language==“English”,], family = Gamma(link = “identity”), control=glmerControl(optimizer = ‘bobyqa’))

## Footnotes

1 Note that in the current study, “ordinal position” refers to the position of the trial within a same-language run, which is different from the same term defined in Meuter & Allport (1999).

2 There were in total 152 switch trials within a list. Therefore, eight out of the 40 stimuli ended up occurring three times instead of four on the switch trials, leaving out each run length once in each language (see Appendix A for more details).

3 EMG was measured to track the time course of speech and to monitor for speech artifacts, but was not analyzed in the current study.

4 A long segment was chosen to provide the baseline for response-locked analysis, which was not used in the current study.

## References

Allport, A., & Wylie, G. (1999). Task-switching: Positive and negative priming of task set. In Attention, Space, and Action: Studies in cognitive neuroscience (pp. 273–296).

Barr, D. J., Levy, R., Scheepers, C., & Tily, H. J. (2013). Random effects structure for confirmatory hypothesis testing: Keep it maximal. Journal of Memory and Language, 68, 255–278. https://doi.org/10.1016/j.jml.2012.11.001

Boersma, P., & Weenink, D. (2016). Praat: doing phonetics by computer [Computer program]. Version 6.0.12. Retrieved 24 January 2016 from http://www.praat.org/.

Christoffels, I. K., Firk, C., & Schiller, N. O. (2007). Bilingual language control: An event-related brain potential study. Brain Research, 1147, 192–208. https://doi.org/10.1016/j.brainres.2007.01.137

Costa, A., & Santesteban, M. (2004). Lexical access in bilingual speech production: Evidence from language switching in highly proficient bilinguals and L2 learners. Journal of Memory and Language, 50, 491–511. https://doi.org/10.1016/j.jml.2004.02.002

Declerck, M., & Philipp, A. M. (2015). A review of control processes and their locus in language switching. Psychonomic Bulletin & Review, 22, 1630–1645. https://doi.org/10.3758/s13423-015-0836-1

Declerck, M., Stephan, D. N., Koch, I., & Philipp, A. M. (2015). The other modality: Auditory stimuli in language switching. Journal of Cognitive Psychology, 27, 685–691. https://doi.org/10.1080/20445911.2015.1026265

Falkenstein, M., Hoormann, J., & Hohnsbein, J. (1999). ERP components in Go/Nogo tasks and their relation to inhibition. Acta Psychologica, 101, 267–291. https://doi.org/10.1016/S0001-6918(99)00008-6

Folstein, J. R., & Van Petten, C. (2008). Influence of cognitive control and mismatch on the N2 component of the ERP: A review. Psychophysiology, 45, 152–170. https://doi.org/10.1111/j.1469-8986.2007.00602.x

Gollan, T. H., Kleinman, D., & Wierenga, C. E. (2014). What’s easier : Doing what you want, or being told what to do ? Cued versus voluntary language and task switching. Journal of Experimental Psychology: General, 143, 2167–2195. https://doi.org/10.1037/a0038006

Green, D. W. (1998). Mental control of the bilingual lexico-semantic system. Bilingualism: Language and Cognition, 1, 67–81. https://doi.org/10.1017/S1366728998000133

Jackson, G. M., Swainson, R., Cunnington, R., & Jackson, S. R. (2001). ERP correlates of executive control during repeated language switching. Bilingualism: Language and Cognition, 4, 169–178. https://doi.org/10.1017/S1366728901000268

Jodo, E., & Kayama, Y. (1992). Relation ofa negative ERP component to response inhibition in a Go/No-go task. Clinical Neurophysiology, 82, 477–482. https://doi.org/10.1016/0013-4694(92)90054-L

Kleinman, D., & Gollan, T. H. (2018). Inhibition accumulates over time at multiple processing levels in bilingual language control. Cognition, 173, 115–132. https://doi.org/10.1016/j.cognition.2018.01.009

Lemhöfer, K., & Broersma, M. (2012). Introducing LexTALE: a quick and valid lexical test for advanced learners of English. Behavior Research Methods, 44, 325–343. https://doi.org/10.3758/s13428-011-0146-0

Lo, S., & Andrews, S. (2015). To transform or not to transform: using generalized linear mixed models to analyse reaction time data. Frontiers in Psychology, 6, 1–16. https://doi.org/10.3389/fpsyg.2015.01171

Maris, E., & Oostenveld, R. (2007). Nonparametric statistical testing of EEG- and MEG-data. Journal of Neuroscience Methods, 164, 177–190. https://doi.org/10.1016/j.jneumeth.2007.03.024

Mayr, U., & Kliegl, R. (2003). Differential effects of cue changes and task changes on task-set selection costs. Journal of Experimental Psychology: Learning, Memory, and Cognition, 29, 362–372. https://doi.org/10.1037/0278-7393.29.3.362

Meuter, R. F. I., & Allport, A. (1999). Bilingual language switching in naming: Asymmetrical costs of language selection. Journal of Memory and Language, 40, 25–40. https://doi.org/10.1006/jmla.1998.2602

Miyake, A., Friedman, N. P., Emerson, M. J., Witzki, A. H., Howerter, A., & Wager, T. D. (2000). The unity and diversity of executive functions and their contributions to complex “Frontal Lobe” tasks: a latent variable analysis. Cognitive Psychology, 41, 49–100. https://doi.org/10.1006/cogp.1999.0734

Oostenveld, R., Fries, P., Maris, E., & Schoffelen, J. M. (2011). FieldTrip: Open source software for advanced analysis of MEG, EEG, and invasive electrophysiological data. Computational Intelligence and Neuroscience, 2011, 1–9. https://doi.org/10.1155/2011/156869

Shao, Z., Roelofs, A., Acheson, D. J., & Meyer, A. S. (2014). Electrophysiological evidence that inhibition supports lexical selection in picture naming. Brain Research, 1586, 130–142. https://doi.org/10.1016/j.brainres.2014.07.009

Sikora, K., Roelofs, A., & Hermans, D. (2016). Electrophysiology of executive control in spoken noun-phrase production: Dynamics of updating, inhibiting, and shifting. Neuropsychologia, 84, 44–53. https://doi.org/10.1016/j.neuropsychologia.2016.01.037

van Casteren, M., & Davis, M. H. (2006). Mix, a program for pseudorandomization. Behavior Research Methods, 38, 584–589. https://doi.org/10.3758/BF03193889

Verhoef, K. M. W., Roelofs, A., & Chwilla, D. J. (2009). Role of inhibition in language switching: Evidence from event-related brain potentials in overt picture naming. Cognition, 110, 84–99. https://doi.org/10.1016/j.cognition.2008.10.013

Verhoef, K. M. W., Roelofs, A., & Chwilla, D. J. (2010). Electrophysiological evidence for endogenous control of attention in switching between languages in overt picture naming. Journal of Cognitive Neuroscience, 22, 1832–1843. https://doi.org/10.1162/jocn.2009.21291

Zheng, X., Roelofs, A., & Lemhöfer, K. (2018). Language selection errors in switching: language priming or cognitive control? Language, Cognition and Neuroscience, 33, 139–147. https://doi.org/10.1080/23273798.2017.1363401

